# The influence of genetic structure on phenotypic diversity in the Australian mango (*Mangifera indica*) gene pool

**DOI:** 10.1101/2022.09.04.506567

**Authors:** Melanie J. Wilkinson, Risa Yamashita, Maddie E. James, Ian S.E. Bally, Natalie L. Dillon, Asjad Ali, Craig M. Hardner, Daniel Ortiz-Barrientos

## Abstract

Genomic selection is a promising breeding technique for tree crops to accelerate the development of new cultivars. However, factors such as genetic structure can create spurious associations between genotype and phenotype due to the shared history between populations with different trait values. Genetic structure can therefore reduce the accuracy of the genotype to phenotype map, a fundamental requirement of genomic selection models. Here, we employed 272 single nucleotide polymorphisms from 208 *Mangifera indica* cultivars to explore whether the genetic structure of the Australian mango gene pool explained variation in tree size, fruit blush colour and intensity. Our results show that genetic structure is weak, but cultivars imported from Southeast Asia (mainly those from Thailand) were genetically differentiated across multiple population genetic analyses. We find that genetic structure was strongly associated with phenotypic diversity in *M. indica*, suggesting that the history of these cultivars could drive spurious associations between loci and key mango phenotypes in the Australian mango gene pool. Incorporating such genetic structure in associations between genotype and phenotype has the potential to improve the accuracy of genomic selection, which can assist the development of new cultivars.

## Introduction

Horticultural tree crops are vital for sustainable food production^1^ and for ornamental and industrial use. Tree crops can be more sustainably cultivated over time than annual field crops, thus helping to manage food supply for an increasing world population^2^. To create new tree fruit cultivars with improved productivity and quality, we must develop breeding technologies that overcome biological limitations to their production.

Tropical species, such as mango, are often large and vigorous^3^, leading to canopies that rapidly outgrow their orchard space. In turn this generates shade, providing a breeding ground for disease^4^. To avoid the adverse effects of tree size, trees are traditionally planted at low density and heavily pruned each year^4^, leading to a reduction in overall production per hectare and an increased cost per unit output. Consequently, a quest to breed smaller, less vigorous trees while maintaining high yields of quality fruit is underway^5,6^. Such efforts will produce mango that can be grown in intensive, high-density orchards that produce more fruit per hectare^7^.

Traditional tree breeding is slow, as evaluations require an assessment of phenotypic performance in mature trees over many years to account for the effects of variable spatial and temporal environments on phenotypic diversity. These evaluations, in combination with a long juvenile phase (typically 2 to 4 years^4^), can result in a selection process of up to or longer than ten years from field planting^8^, making the rapid development of new cultivars unfeasible. The time for cultivar development could be reduced by predicting future phenotypic performance in young individuals using genomic selection, as demonstrated in apples^9^, sweet cherry^10^ and strawberry^11^. Genomic selection uses genotype to phenotype maps from a training population to predict phenotypic variation in untested populations using marker data^12,13^. Thus, once a genomic selection model has been created, the length and expense of phenotyping key traits may be reduced. Genomic selection for tree size and vigour of progeny could therefore improve the breeding process and reduce the cost of mango breeding compared to traditional breeding approaches.

The primary assumption of genomic selection is that genetic markers are in close physical linkage on a chromosome with the causative loci that contribute to the trait of interest^14^. In general, the closer the marker is to the causative loci, the more accurate the genotype to phenotype map. However, genetic structure can create statistical associations between loci that are not physically linked. This occurs because evolutionary forces such as migration, drift and mutation can make allelic combinations between unlinked loci more common than expected by chance^15^. Genetic structure can therefore create spurious associations between genetic markers and traits. Furthermore, genetic structure is often prevalent in modern crops, particularly those moving across the world via human migrations, which likely experienced drastic fluctuations in population size, and suffered from inbreeding after crossing genetically related individuals with favourable traits^16^.

Differentiating uninformative loci due to genetic structure from those that are linked to causative loci is a common problem observed in genetic studies of human disease^17,18^ and in the study of trait evolution across diverse taxa^19-23^. Fortunately, we can improve the accuracy of the genotype to phenotype map by accounting for genetic covariation between traits and markers due to genetic structure^24-26^, a practice that can potentially improve the quality of horticultural breeding programs that start from highly variable germplasm collections. Here, we evaluate the assumption that horticultural trait variation segregates independently from genetic structure using *Mangifera indica* in the gene pool of the Australian Mango Breeding Program.

Mango is a major horticultural tree crop worldwide, yet an understanding of the domestication history is still debated. The centre of origin of the genus *Mangifera* is Southeast Asia, but the origin of the species *M. indica* is still under question. Based on the fossil record, Mukherjee^27^ and Blume^28^ suggested mango originated in the Malay Archipelago less than 2.58 million years ago. However, recent molecular taxonomy suggests it evolved within a large area of Northwest Myanmar, Bangladesh and Northeast India^29^. From this area, human migration and trading led to the dispersion of mangoes to many regions of the world^30^.

Several studies have evaluated the genetic structure of domesticated mango^31-38^, yet to our knowledge there have been no published studies on the effects of genetic structure on phenotypic variation in mango cultivars. One study with 60 mango cultivars from India accounted for genetic structure in a marker trait analysis^35^, however, Lal et al.^35^ did not assess the effect of genetic structure on their genotype to phenotype map. Without understanding the effect of genetic structure on phenotypic diversity, we do not know whether we are creating false associations between genetic markers and key mango traits. Here, we directly examined the effects of genetic structure on the creation of spurious associations between genetic markers and three traits – tree size, blush colour and intensity – in the Australian mango gene pool. We assessed 272 SNP markers genotyped in 208 *M. indica* landraces imported worldwide and reveal statistical associations between genetic marker and trait arising from genetic structure. We discuss the potential impact of these results on the accuracy of genomic selection on young trees.

## Results

### Genetic structure in the Australian mango gene pool

Genetic structure was found in both a hierarchical cluster analysis (HCA) and a principal component analysis (PCA) across all 208 *M. indica* cultivars (Fig. 1). Consistent with a recent origin of all cultivars, the HCA created a dendrogram with only short branches in the centre (Fig. 1a), indicating few genetic differences separate the clusters. The optimal number of genetic clusters was K=4 as indicated by the HCA, the elbow plot, and the STRUCTURE analysis (explained below). The elbow plot from the HCA shows diminishing returns in the amount of variance explained after five clusters (Fig. S1). In the dendrogram, cluster 1 is the most genetically differentiated cluster, which only contains cultivars imported from Southeast Asia. Cluster 1 is most distinct from clusters 2 and 3 while cluster 4 is more similar to cluster 1 (Fig. 1a), and contains a mixture of samples across geographical regions (e.g., South Asia, Southeast Asia, Americas and Oceania; Table 1; Fig. 2). In the reduced PC space (Fig. 1b), genetic clusters largely overlap, with South Asian cultivars (mostly Indian cultivars) primarily concentrated in the centre of the multivariate space. Genetic clusters from Southeast Asia, the Americas, and Oceania occur towards the edges of the genotypic space, with Southeast Asia distinctly separated in the PC1 axis.

**Table 1.** The number of cultivars of *M. indica* from each country of import and their assigned genetic clusters from the hierarchical cluster analysis for K=4 calculated from 272 biallelic SNPs. Countries have been grouped into six geographical regions of import.

| Country | Country code | cluster 1 | cluster 2 | cluster 3 | cluster 4 | Country Total |
| --- | --- | --- | --- | --- | --- | --- |
| **Africa** |  |  |  |  |  |  |
| East Africa | EAF |  |  | 1 |  | 1 |
| Kenya | KEN |  | 1 |  |  | 1 |
| South Africa | ZAF |  | 3 |  |  | 3 |
| **Americas** |  |  |  |  |  |  |
| Brazil | BRA |  | 1 | 1 |  | 2 |
| Jamaica | JAM |  | 1 | 1 | 1 | 3 |
| Saint Lucia | LCA |  |  | 1 |  | 1 |
| United States of America | USA |  | 32 | 8 |  | 40 |
| **Middle East** |  |  |  |  |  |  |
| Israel | ISR |  | 3 | 1 |  | 4 |
| **Oceania** |  |  |  |  |  |  |
| Australia | AUS |  | 37 | 15 |  | 52 |
| French Polynesia | PYF |  |  | 1 | 1 | 2 |
| **South Asia** |  |  |  |  |  |  |
| India | IND |  | 14 | 15 | 5 | 34 |
| Pakistan | PAK |  | 1 |  |  | 1 |
| Sri Lanka | LKA |  | 1 | 2 |  | 3 |
| **Southeast Asia** |  |  |  |  |  |  |
| Indonesia | IDN |  | 2 | 6 | 2 | 10 |
| Malaysia | MYS | 1 |  | 2 | 1 | 4 |
| Malesia | MLS |  | 1 | 2 | 1 | 4 |
| Myanmar | MMR | 1 |  |  |  | 1 |
| Philippines | PHL | 3 |  |  |  | 3 |
| Singapore | SGP |  |  | 1 |  | 1 |
| Thailand | THA | 18 | 1 | 3 |  | 22 |
| Vietnam | VNM | 5 |  | 4 |  | 9 |
| unknown |  |  | 4 | 3 |  | 7 |
| **Cluster total** |  | 28 | 102 | 67 | 11 | 208 |

**Fig. 1.**
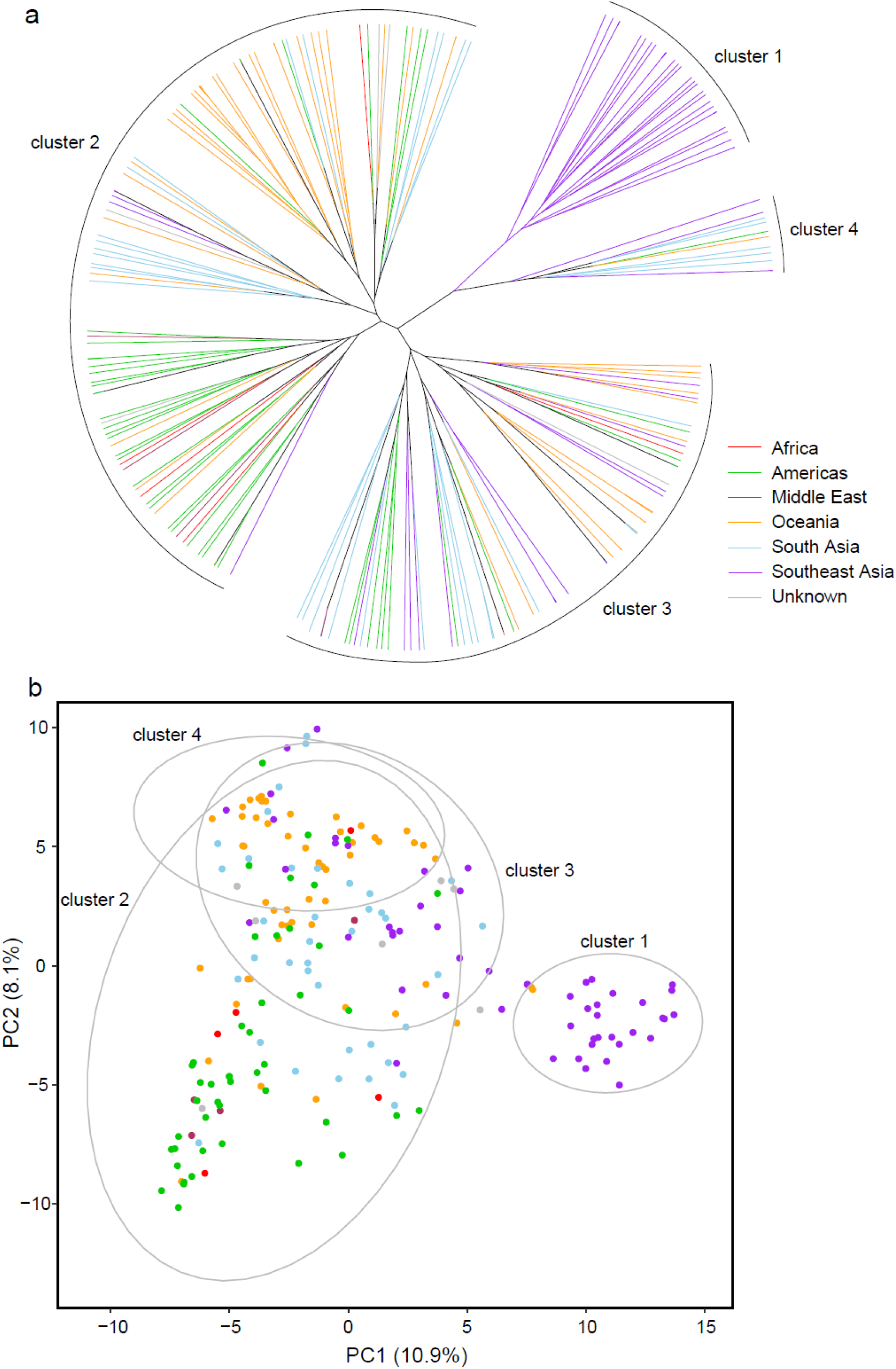
Genetic structure analyses for K=4 of the 208 cultivars of *M. indica* from six geographical regions across the world. **a)** A circular dendrogram showing the hierarchical cluster analysis using complete linkage clustering. Each branch represents an individual with the colour of the branch representing the geographical region the sample was imported into Australia from. **b)** Principal components analysis, where the ellipses (95% probability) represent the four clusters from the hierarchical cluster analysis.

**Fig. 2.**
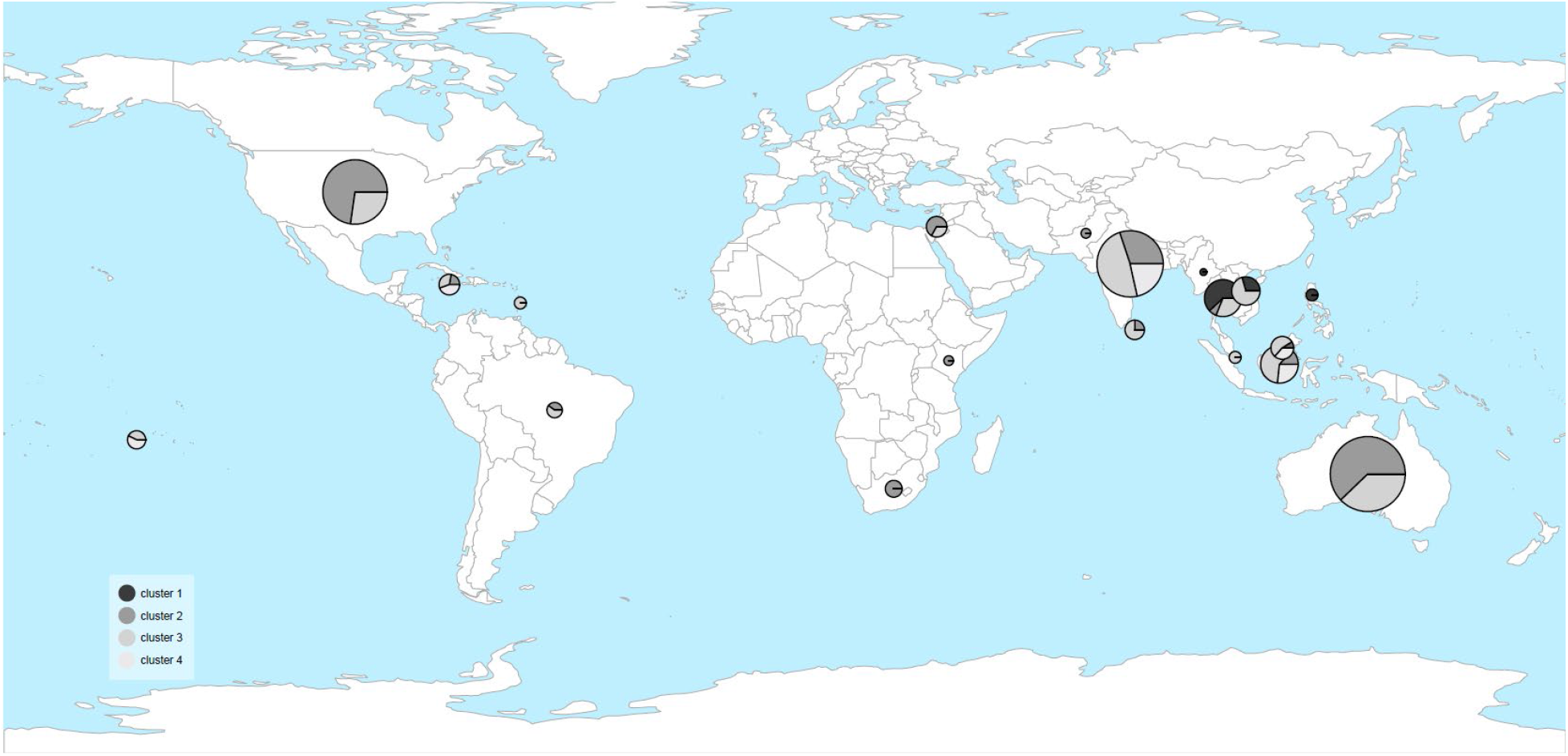
Genetic structure across geography of the 208 *M. indica* cultivars. Cluster numbers (K=4) were determined using a hierarchical cluster analysis (Fig. 1). The size of each pie chart reflects the number of cultivars imported from each country.

In agreement with the HCA and PCA results above, we identified genetic clusters across the 208 *M. indica* cultivars (Fig. 3) using the Bayesian clustering approach implemented in STRUCTURE^39^. Most cultivars contain large amounts of admixture or shared ancestral polymorphism, where portions of their genome were assigned to different genetic groups. When genetic differentiation was separated into only two groups (K=2, see Methods), Southeast Asia formed one group, while all other cultivars were in a second group (Fig. 3).

**Fig. 3.**
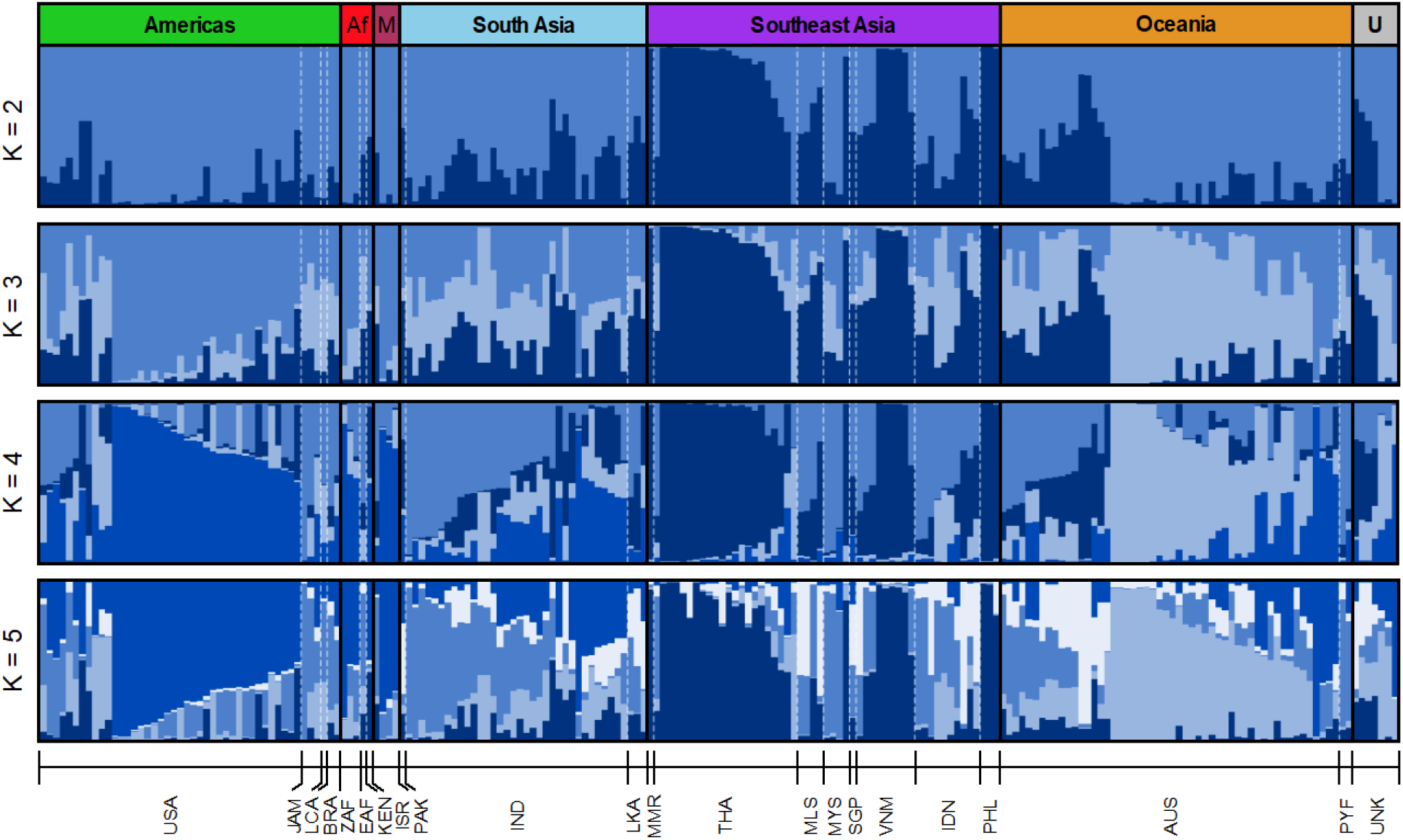
Genetic structure of 208 *M. indica* individuals using STRUCTURE for K=2-5. Each bar represents an individual with the shades of blue representing the ancestry proportions to each cluster. Individuals are sorted by geographical region (black lines) and countries (white dotted lines). Country codes are listed at the bottom.

Relaxing this constraint to K=3 revealed the Americas and Oceania cultivars each form their own group. Populations are almost indistinguishable when K is larger than 4. Consistent with the elbow plot discussed above, the Evanno method^40^ and the log probability of K values show K=4 was the optimal number of clusters (Fig. S2). Most cultivars show signatures of admixture as indicated by diversity from multiple groups. Admixture signals are particularly pronounced in cultivars from South Asia, mainly those from India, which do not form a distinct genetic group with any K-value.

Together, the HCA, PCA and STRUCTURE results suggest that mango landraces of the Australian mango gene pool consist of four genetic groups. Southeast Asian cultivars are most differentiated relative to the rest of the world, suggesting that these cultivars might have evolved differently, thus creating a heterogenous gene pool for cultivar creation in the Australian Mango Breeding Program.

### Patterns of genetic diversity across the Australian mango gene pool

Genetic diversity analyses revealed high levels of heterozygosity and variable patterns of inbreeding across regions (Table 2). Levels of expected heterozygosity (H^E^) and observed heterozygosity (H_O_) were high across all cultivar regions, with the Americas having the highest levels of observed heterozygosity (H_O_ = 0.49) and Southeast Asia having the lowest (H_O_ = 0.39). Cultivars from the Americas contain an excess of heterozygote individuals (i.e., a negative inbreeding coefficient; F_IS_ = -0.11; 95% CI, -0.13 to -0.08). On the other hand, cultivars from Southeast Asia are mildly inbred (i.e., a positive inbreeding coefficient; F_IS_ = 0.08; 95% CI, 0.06 to 0.11). Private alleles were absent in all regions, indicating either a large intermixing population or high levels of ancestral polymorphism that has not been sorted across geographic regions.

**Table 2.** Genetic diversity for 208 *M. indica* cultivars across six geographic regions using 272 SNPs.

| Region | H<sub>O</sub> | H<sub>E</sub> | F<sub>IS</sub> (95% CI's) | Pr |
| --- | --- | --- | --- | --- |
| Africa | 0.46 | 0.45 | -0.02 (-0.08 – 0.05) | 0 |
| Americas | 0.49 | 0.44 | -0.11* (-0.13 – -0.08) | 0 |
| Middle East | 0.46 | 0.45 | -0.01 (-0.08 – 0.05) | 0 |
| Oceania | 0.43 | 0.42 | -0.02 (-0.05 – 0.00) | 0 |
| South Asia | 0.45 | 0.45 | 0.00 (-0.03 – 0.02) | 0 |
| Southeast Asia | 0.39 | 0.42 | 0.08* (0.06 – 0.11) | 0 |

H<sub>O</sub>, observed heterozygosity; H<sub>E</sub>, expected heterozygosity; F<sub>IS</sub>, inbreeding co-efficient with 95% confidence intervals, where \*CI's do not overlap with 0; Pr, the number of private alleles.

Genetic differentiation comparisons showed variable patterns of F_ST_ between genetic clusters and between regions of import. Comparisons between regions have low levels of F_ST_, which range from -0.016 to 0.112 (Table 3a). Southeast Asia and the Middle East, closely followed by the comparison between Southeast Asia and the Americas, showed the highest level of genetic differentiation (F_ST_ = 0.112 and 0.107, respectively). In contrast, F_ST_ between clusters ranged from 0.051 to 0.286 with cluster 1 comparisons having the highest values (Table 3b). Overall, there is low genetic divergence amongst regions of the Australian mango gene pool and high genetic divergence between genetic clusters.

**Table 3.** Pairwise F_ST_ for *M. indica*. **a)** F_ST_ estimates for the geographical regions and **b)** clusters. F_ST_ estimates are below the diagonal, 95% confidence interval above the diagonal are based on 1,000 bootstrap replicates. Clusters (K=4) were identified using hierarchical cluster analysis (Fig. 1).

| Region | Africa | Americas | Middle East | Oceania | South Asia | Southeast Asia |
| --- | --- | --- | --- | --- | --- | --- |
| **Africa** | - | -0.012 – 0.004 | -0.036 – 0.005 | 0.022 – 0.055 | 0.010 – 0.035 | 0.062 – 0.095 |
| **Americas** | -0.004 | - | -0.008 – 0.012 | 0.051 – 0.07 | 0.036 – 0.053 | 0.092 – 0.121 |
| **Middle East** | -0.016 | 0.002 | - | 0.059 – 0.099 | 0.009 – 0.036 | 0.088 – 0.134 |
| **Oceania** | 0.039* | 0.060* | 0.079* | - | 0.031 – 0.046 | 0.070 – 0.095 |
| **South Asia** | 0.022* | 0.044* | 0.022* | 0.038* | - | 0.049 – 0.069 |
| **Southeast Asia** | 0.078* | 0.107* | 0.112* | 0.082* | 0.060* | - |

**b**
| Cluster | 1 | 2 | 3 | 4 |
| --- | --- | --- | --- | --- |
| **1** | - | 0.191 – 0.151 | 0.174 – 0.135 | 0.322 – 0.250 |
| **2** | 0.171* | - | 0.059 – 0.043 | 0.170 – 0.127 |
| **3** | 0.153* | 0.051* | - | 0.074 – 0.05 |
| **4** | 0.286* | 0.148* | 0.062* | - |

\* 95% confidence intervals do not overlap with 0.

### Genetic structure and region of import influence phenotypic diversity

Phenotypic correlation analyses revealed associations between fruit traits but not between them and trunk circumference. Trunk circumference, a continuous trait, was highly variable at nine years, ranging from 27-70cm, while categorical fruit traits were less variable (see Fig. S3 for photos of each fruit blush colour and intensity category). In a single-factor linear model, fruit blush colour and intensity were strongly correlated (LR χ^2^=373.168, df=4, p<0.0001, R^2^=0.61). However, given that 39% of mango cultivars lacked fruit blush colour, and therefore lacked fruit blush intensity, we removed ‘no blush’ and retested the association. It led to a significant yet weaker association between the fruit traits (LR χ^2^=95.077, df=3, p<0.0001, R^2^=0.28). We found no correlation between trunk circumference and fruit blush colour (Fig. S4; F_4,203_=1.093, p=0.3613, R^2^=0.02) and trunk circumference and fruit blush intensity (Fig. S5; F_4,203_=1.473, p=0.2118, R^2^=0.03), suggesting trunk circumference is likely to be genetically independent of these fruit traits.

Fruit blush traits are strongly associated with the region of import in the Australian mango gene pool. In single trait linear models, region of import showed a significant effect on fruit blush colour (Fig. 4a; LR χ^2^=77.768, df=12, p<0.0001, R^2^=0.14) and fruit blush intensity (Fig. 4b; LR χ^2^=98.936, df=3, p<0.0001, R^2^=0.18), but not trunk circumference (F_3,188_=, p=0.1200, R^2^=0.03). Of the regions that had more than ten samples, trunk circumference ranged from a mean of 48.1±1.8 (n=38) in South Asia to a mean of 52.5±1.3 (n=46) in the Americas (Table S1). For fruit blush colour (Table S2), 67% of cultivars from Southeast Asia had no blush colour, while only 11% from the Americas had no blush, with most having red blush (43%). For fruit blush intensity (Table S3), the Americas had 41% of cultivars with a medium blush intensity that resembled the Haden cultivar. In comparison, Oceania had 39% of cultivars with slight blush intensity resembling the Kensington Pride cultivar. Contrastingly, 94% of Southeast Asian cultivars and 82% of South Asian cultivars had no blush or barely visible blush intensity.

**Fig. 4.**
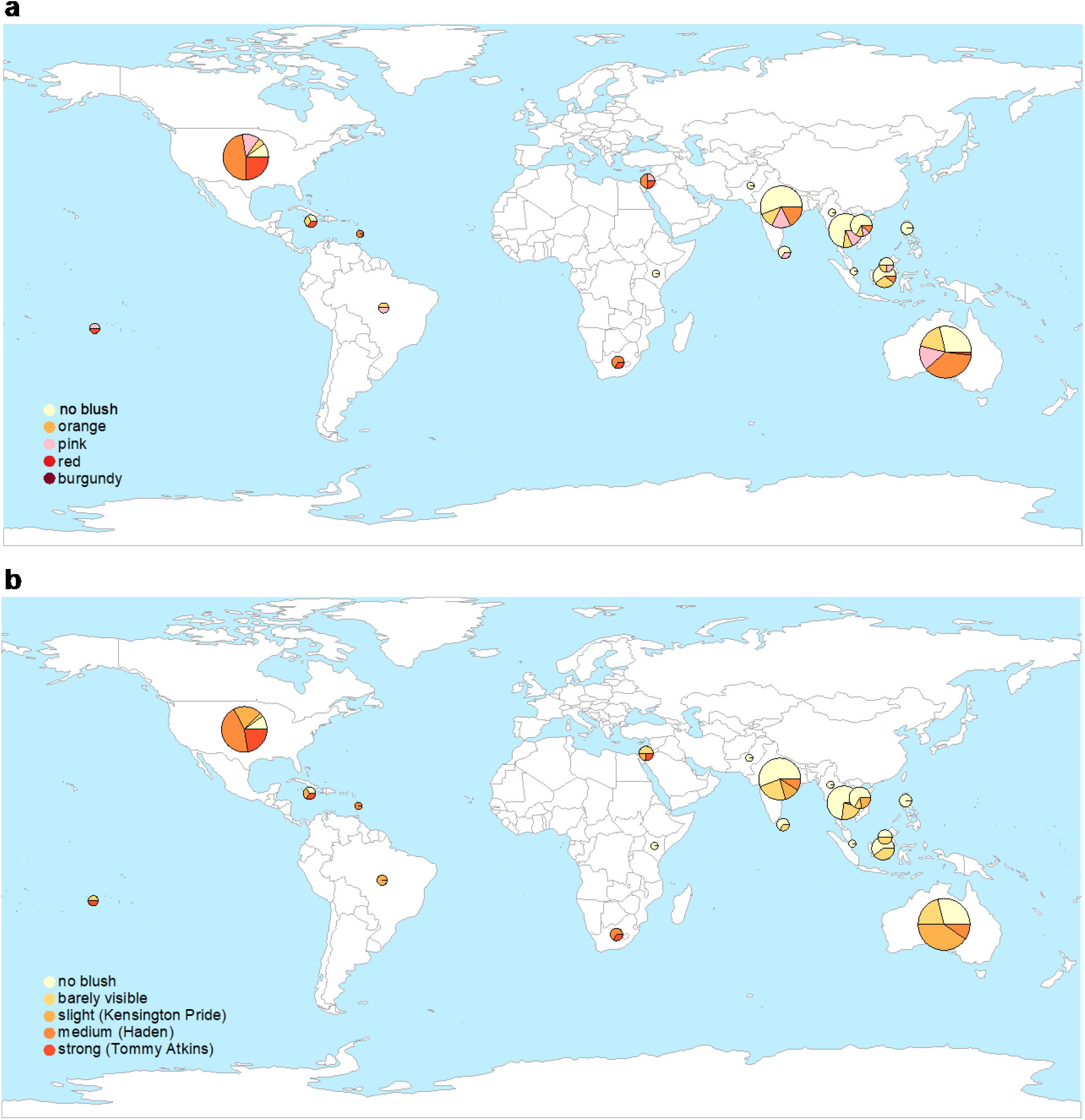
Fruit blush colour and intensity across geography of the 208 *M. indica* cultivars. **a)** Fruit blush colour is split into five categories. **b)** Fruit blush intensity increases from no blush to strong blush on an ordinal scale, where the cultivars in brackets best reflect the colour intensity. The size of each pie chart reflects the number of cultivars imported from each country.

Phenotypic diversity is associated with the four clusters assigned in the HCA. Cluster assignment had a significant effect on the presence of blush (LR χ^2^=28.046, df=3, p<0001, R^2^=0.10), where 18% of individuals in cluster 1 had blush, whereas 70% and 69% of individuals from clusters 2 and 3 had blush, respectively. Cluster 1 is more likely to have lower blush intensity than the other clusters when the ‘no blush’ category is excluded (LR χ^2^=12.274, df=3, p=0.0065, R^2^=0.04; odds ratios between cluster 1 and clusters 2-4 ranged from 3.8-10.5). Cluster also had a significant effect on trunk circumference (F_3,204_=18.410, p<0001, R^2^=0.21), where cluster 1 (mean=52.3±1.5, n=28) and cluster 2 (mean=53.7±0.8, n=102) had the largest trunk circumference and cluster 4 had the smallest (mean=36.8±2.9, n=11). Overall, we expect that genetic diversity and factors specific to the region of import will likely influence the genotype to phenotype map of these key mango traits.

## Discussion

Genetic structure arises from evolutionary processes such as mutation, migration and genetic drift, which drive shifts in allelic frequency that could cause statistical associations between random genetic markers and traits^41^. Such variation arising from genetic structure is often confounded with loci contributing to trait variation in association studies^17-23^ and can misrepresent the genotype to phenotype map assumed in genomic selection models. Our study shows how genetic structure in *M. indica* can lead to statistical associations between genetic markers and three phenotypic traits measured in this study – trunk circumference, fruit blush colour and intensity. This suggests that the genetic architecture of these horticultural traits contains noise arising from the conflation of phenotypic and historical differences in the Australian mango gene pool. Such noise can create spurious associations that hinder the selection of new cultivars.

Genetic variability and divergence in the Australian mango gene pool can be understood in two ways. On one hand, cultivars imported from different regions are weakly differentiated. On the other hand, genetic clusters are strongly differentiated, implying the existence of clear genetic groups across the world. Results described in Fig. 2 reveal that genetic clusters are distributed across regions, implying that their genetic structure is shared amongst geographic regions. The net effect of this nested relationship between geographic region and genetic cluster is low F_ST_ values amongst the regions, yet high levels of F_ST_ amongst genetic clusters. This relationship can be used to hypothesise the causes of genetic divergence in the Australian mango gene pool.

In our study, Cluster 1 (containing only Southeast Asian countries) comprises the most genetically differentiated cultivars from across the world. Previous studies support this observation; Warschefsky and von Wettberg^31^ showed that cultivars from Southeast Asia cluster together in a STRUCTURE plot, and Dillon et al.^32^ reached a similar conclusion using genetic distance analyses of 254 mango cultivars. Cultural differences amongst regions might help explain some of the variations in phenotype between Southeast Asia and the rest of the world. For example, Southeast Asia makes savoury dishes with mangoes, which might have led to selection of immature mangoes that stay green while ripening and therefore lack blush^31^. On the other hand, red blush is favoured around the world^42^, likely accentuating some of the genetic differentiation between cultivars from Southeast Asia and the rest of the world.

Artificial selection for these cultural preferences may have driven some of the population differentiation identified in the Australian mango gene pool. It is well accepted that selecting for one trait can incidentally lead to the evolution of other traits^43,44^. The genetic architecture of selected traits will largely determine the extent of this correlated evolution. In this study, we show that fruit blush traits are highly correlated, which might imply a shared genetic architecture, thus selection for either of these traits would at least partially drive the evolution of the other. For example, if low levels of blush have been preferentially selected in Southeast Asia, then this would have likely driven the evolution of low blush intensity, but not trunk circumference that has a unique genetic architecture. Another aspect of the genetic architecture that will likely drive genetic differentiation across the genome is selection for polygenic traits and pleiotropic genes. Trunk circumference is a polygenic trait^45^ and therefore selection for this trait may lead to changes in allelic frequencies across many loci. In contrast, fruit colour pigments and their levels are often controlled by fewer loci in simpler biochemical pathways^46-48^ relative to those controlling plant growth and development^49-51^. Future studies should consider the influence of the genetic architecture of selected traits in driving genetic differentiation.

Polyembryony could have contributed to some of the genetic differences between Southeast Asia and other cultivars. Southeast Asian cultivars are typically polyembryonic where all but one (the zygotic embryo) of the multiple somatic embryos are genetically identical to the maternal parent. Polyembryony is likely to easily be maintained under moderate to strong selection as it is thought to be inherited through a single dominant gene^52,53^. A high level of polyembryony can freeze the genetic diversity in a population, as instead of allowing hybridisation and the creation of unique individuals through recombination, it propagates genetically identical individuals^54^. Polyembryony can therefore create genetic bottlenecks if only a fraction of the original genetic diversity is propagated, which is consistent with the signature of inbreeding in Southeast Asian cultivars found in this study. Furthermore, previous studies have found genetic clustering of mango cultivars according to their ability to produce polyembryonic seed^55,56^. However, embryo type is conflated with geographic region in these studies, where Southeast Asia cultivars dominate the polyembryony types. Therefore, without future work teasing apart the contribution of polyembryony and geographic region than we lack an understanding of the various causes of polyembryonic selection and inbreeding on the genetic diversity of tree crops.

Genetic diversity and the partitioning of genetic structure influences prediction accuracy in genomic selection models across horticultural crops^16,24,26,57-59^. For instance, increasing genetic diversity by using a variety of different races or genetic clusters in the training and validation sets produced higher prediction accuracies in rice^58^, sorghum^16^ and wheat^59^. But genetic diversity is known to reduce prediction accuracy when estimation error is high, which occurs in small populations or when there is low marker density^59,60^. By definition, using markers close to the causative variants will augment prediction accuracy during breeding^61^; however, this is hard to achieve with low-density genotyping techniques such as SNP chips and Genotyping by Sequencing. With sequencing that covers the entire genome (e.g., whole genome sequencing), factors that are influenced by the effect of linkage disequilibrium can be better controlled, such as finding markers in tight linkage with causative loci. As such, population size, marker density, genetic structure of the population, and the genetic architecture of the chosen traits will play a significant role in the accuracy of genomic selection models.

To ameliorate the adverse effects of genetic structure in genomic selection models, there are two major approaches used across horticultural crops^16,24,26,57,59^. The first approach includes principal components from genetic structure analyses as covariates in the model^62-65^. However, this method can double-count genetic structure because some of the elements are included in the model through the genomic relationship matrix^66^. Another common approach for accounting for genetic structure in genomic selection models is ensuring an equal contribution across genetic clusters in both training and validation sets. This stratified sampling approach has been shown to increase prediction accuracy in sorghum^16^ and maize^67^, and could be an effective method in the Australian mango gene pool. In general, choosing the most accurate genomic selection model will largely depend on the breeding population’s characteristics and the population’s genetic structure and size.

## Conclusion

The results of this study reveal that a horticultural species spread across the world has a genetic structure that can create statistical associations between key traits and genetic markers. To remove the effects of spurious markers, breeders should fully characterise the genetic structure of their breeding population. This will allow them to incorporate sample stratification to improve the performance of genomic selection models. Together with best practices of genomic selection (e.g., whole genome sequencing and large population size), these considerations can improve the genotype to phenotype map to assist choosing individuals with accurate breeding values and help advance future parental selection. We hope our study encourages other horticultural breeding programs to follow similar methods.

## Methods

### Cultivars

A total of 208 *M. indica* landraces were used from the gene pool collection of the Australian Mango Breeding Program managed by the Department of Agriculture and Fisheries at Walkamin Research Station, Queensland. Although all cultivars were grown in Australia, they were imported from 21 countries across six geographical regions, with most samples originating from Australia (n=52), the United States of America (n=40), India (n=34), and Thailand (n=22). See Table 1 for the complete list of countries and sample sizes.

### Genotyping

To identify some of the genotypic diversity in the Australian mango gene pool, we used the genotypes from Kuhn et al.^68^. DNA isolation for these genotypes was described in Kuhn et al.^69^. Briefly, young leaf samples were collected from Walkamin Research Station and the glasshouse at Mareeba Research Facility, Queensland. DNA was extracted using 20mg of fresh sample with the Qiagen Plant DNeasy kit. SNP genotyping was performed on these DNA samples using the Fluidigm EP-1 platform with 384 biallelic SNP markers. Finally, 272 SNP markers were selected for further analyses, where 236 markers belonged to one of 20 linkage groups (7-20 markers per linkage group), and the remaining 36 markers were unknown^68^. Genotypically identical individuals across the 272 SNPs were consolidated, leaving 208 mango cultivars for the analyses. On average, 98% of the 272 SNPs used in this study were successfully genotyped in every individual.

### Hierarchical Cluster Analysis

To examine the genotypic clustering of the mango cultivars due to genotypic similarity, we performed a hierarchical cluster analysis (HCA) of the 208 *M. indica* cultivars. First, pairwise genetic distances between all cultivars were calculated using the percentage method by the “ape” v.5.3 R-package^70^. The HCA was conducted by “stats” v. 3.6.2 R-package^71^ with complete linkage clustering. This computes all pairwise dissimilarities between the cultivars in a cluster and cultivars in another cluster and considers the largest value of these dissimilarities as a distance between the two clusters. To assess the optimal number of clusters, we used the elbow method^72^, which plots the total within-cluster sum of squares (WSS) against the number of clusters to show the ‘elbow’ where the WSS rate of decrease slows and indicates diminishing returns with more clusters^73^.

### Principal Components Analysis

We assessed the major patterns of genetic similarity among the 208 mango cultivars in multivariate space using a principal components analysis (PCA) with 272 SNPs. Missing SNP data were imputed using the regularised iterative PCA algorithm with the “missMDA” v1.17 R-package^74^. The PCA was performed using the “stats” v3.6.2 R-package^71^. Ellipses were constructed for each of the four clusters in the HCA to identify the position of every individual in a cluster in multivariate space with 95% probability.

### STRUCTURE analysis

We determined levels of admixture between all 208 *M. indica* individuals with STRUCTURE v2.3.4^39^. STRUCTURE is a Bayesian Markov chain Monte Carlo (MCMC) program that assigns individuals into genetic clusters (K) based on their genotypes by assuming Hardy Weinberg equilibrium within a cluster. It gives each individual an admixture coefficient to depict the proportion of the genome originating from a particular K cluster. We ran the admixture model and the correlated allele frequency model^75^ with ten independent runs of 100,000 burn-in and 100,000 MCMC iterations for K=1-7. We visually inspected summary statistics of MCMC runs to ensure convergence of model parameters. Results were summarised and plotted in the “pophelper” v2.2.7 R-package^76^. The optimal K value (which represents the most likely number of sub-populations) was estimated by the Evanno method^40^, which uses the second-order rate of change in the log probability of data between successive K values in the R-package StructureSelector^77^. The optimal K value was also estimated using LnP(K), the mean log probability of the data. We also followed suggestions by Pritchard et al.^78^ and Lawson et al.^79^ and plotted the lowest K values that capture the primary structure in the data.

### Genetic diversity and genetic differentiation

To examine the level of differentiation between the clusters and geographical regions, Weir and Cockerham’s pairwise F_ST_ and 95% confidence intervals were estimated by “hierfstat” v.0.4.22 R-package^80^. Each cultivar was assigned to a cluster based on the HCA, and each country of import was grouped into six geographic regions. We calculated 95% confidence intervals for each pairwise comparison using 1,000 bootstrap replicates. Significance was determined by whether the confidence interval overlapped with 0.

Measures of genetic diversity were calculated for all 208 *M. indica* cultivars for each of the six geographic regions. A genind object was created in “adegenet” v.2.1.2^81,82^ for input into “hierfstat” v.0.4.22^80^ to calculate observed heterozygosity (H_o_), expected heterozygosity (H_E_) and the inbreeding coefficient (F_IS_). To determine whether F_IS_ was significantly different from 0, we calculated 95% confidence intervals for each pairwise comparison using 1,000 bootstrap replicates. The number of private alleles (Pr) was calculated with the “poppr” v.2.8.6 R-package^83,84^.

### Phenotyping

To capture some of the phenotypic diversity in the Australian mango gene pool, we measured three traits in all 208 mango cultivars – trunk circumference, fruit blush colour and fruit blush intensity. Trunk circumference was used as a proxy for tree size, as it has been found to be a strong indicator of tree size in other tree crops^85-87^. Trunk circumference was measured 10cm above the graft when the trees were nine years old at Walkamin Research Station. Fruit blush colour and intensity were assessed using ten ripe fruits from each mango cultivar. Fruit blush included five categories: no blush, orange, pink, red and burgundy (Fig. S3a). Fruit blush intensity was recorded as five ordinal variables increasing in colour intensity (Fig. S3b), where the cultivars in brackets best reflect the colour intensity: no blush, barely visible, slight (Kensington Pride), medium (Haden) and strong (Tommy Atkins).

### The effect of region of import and genetic structure on phenotypic diversity

Tests of association were undertaken to examine the relationship between traits. Chi-square likelihood ratios were used to test phenotypic association amongst the categorical traits of fruit blush and intensity. We then performed the same analysis with the ‘no blush’ category removed to test whether the association remains. A linear model was performed to test for an association between trunk circumference and fruit blush colour, and also trunk circumference and fruit blush intensity.

To understand the effect of region of import on both genotype and phenotype in the Australian mango gene pool, we tested its association with genetic structure and phenotypic diversity. We investigated the influence of geographical region on phenotypic diversity for three key mango phenotypes – trunk circumference, fruit blush colour and intensity. We performed a likelihood-ratio chi-square test for fruit blush colour (categorical) and intensity (ordinal) against the region of import and a linear model for trunk circumference. Region of import was the explanatory variable in each model and included the regions shown in Table 1, excluding unknown regions (n=7) and regions with low samples sizes, including Middle East (n=4), and Africa (n=5).

We then tested for an effect of genetic structure on the three phenotypes using the optimal cluster assignment of K=4 from the HCA. A likelihood-ratio chi-square test was performed for whether the cluster explained the presence (n=127) vs absence of blush (n=81), irrespective of the intensity of blush. We then removed the individuals with no blush from the dataset to test whether there was a significant difference in fruit blush intensity between clusters for just the individuals with fruit blush using a likelihood-ratio chi-square test with an odds ratio. Finally, we performed a mixed linear model to test the effect of cluster on trunk circumference. JMP v15.2.0 (SAS 2015) produced all statistical results reported here.

## Acknowledgements

We thank David Kuhn formerly of Agriculture Research Services, United States Department of Agriculture (ARS-USDA) and Barbara Freeman of ARS-USDA for providing the genotypes. This research was undertaken as part of the National Tree Genomics Program – Phenotype Prediction project (AS17000) which is funded by the Hort Frontiers Advanced Production Systems fund, part of the Hort Frontiers strategic partnership initiative developed by Hort Innovation, with co-investment from The University of Queensland and Queensland Government, and contributions from the Australian Government.

## Author contributions

M.W., C.H. and D.O. designed the study. I.B., N.D and A.A. collected and curated the data. M.W., R.Y. and M.J. performed data analysis. M.W, M.J., R.Y. and D.O. wrote the manuscript. D.O. and C.H. secured funding and were mentors. All authors reviewed the manuscript.

## Data Availability

The datasets generated during and/or analysed during the current study are available from the corresponding author on reasonable request.

