## Supplementary Tables and Figures for "The influence of genetic structure on phenotypic diversity in the Australian mango (*Mangifera indica*) gene pool"

**Table S1. Mean and standard error (SE) for trunk circumference for the six geographic regions of import of *M. indica*.**

| Region | N samples | Mean | SE |
|---|---|---|---|
| Africa | 5 | 53.4 | 6.7 |
| Americas | 46 | 52.5 | 1.3 |
| Middle East | 4 | 55.5 | 1.3 |
| Oceania | 54 | 51.1 | 1.1 |
| South Asia | 38 | 48.1 | 1.8 |
| Southeast Asia | 54 | 49.3 | 1.2 |
| Unknown | 7 | 53.1 | 3.9 |

**Table S2. Fruit blush colour percentages for the six geographic regions of import of *M. indica*.**

| Region | no blush | orange | pink | red | burgundy |
|---|---|---|---|---|---|
| Africa | 20 | 0 | 0 | 60 | 20 |
| Americas | 11 | 9 | 13 | 43 | 24 |
| Middle East | 0 | 0 | 25 | 50 | 25 |
| Oceania | 28 | 17 | 17 | 35 | 4 |
| South Asia | 58 | 11 | 16 | 16 | 0 |
| Southeast Asia | 67 | 15 | 15 | 4 | 0 |
| Unknown | 29 | 14 | 29 | 14 | 14 |

**Table S3. Fruit blush intensity percentages for the six geographic regions of import of *M. indica*.**

| Region | no blush | barely visible | slight (Kensington Pride) | medium (Haden) | strong (Tommy Atkins) |
|---|---|---|---|---|---|
| Africa | 20 | 0 | 20 | 40 | 20 |
| Americas | 11 | 2 | 24 | 41 | 22 |
| Middle East | 0 | 50 | 25 | 0 | 25 |
| Oceania | 28 | 22 | 39 | 9 | 2 |
| South Asia | 58 | 24 | 11 | 8 | 0 |
| Southeast Asia | 67 | 28 | 6 | 0 | 0 |
| Unknown | 29 | 0 | 14 | 29 | 29 |

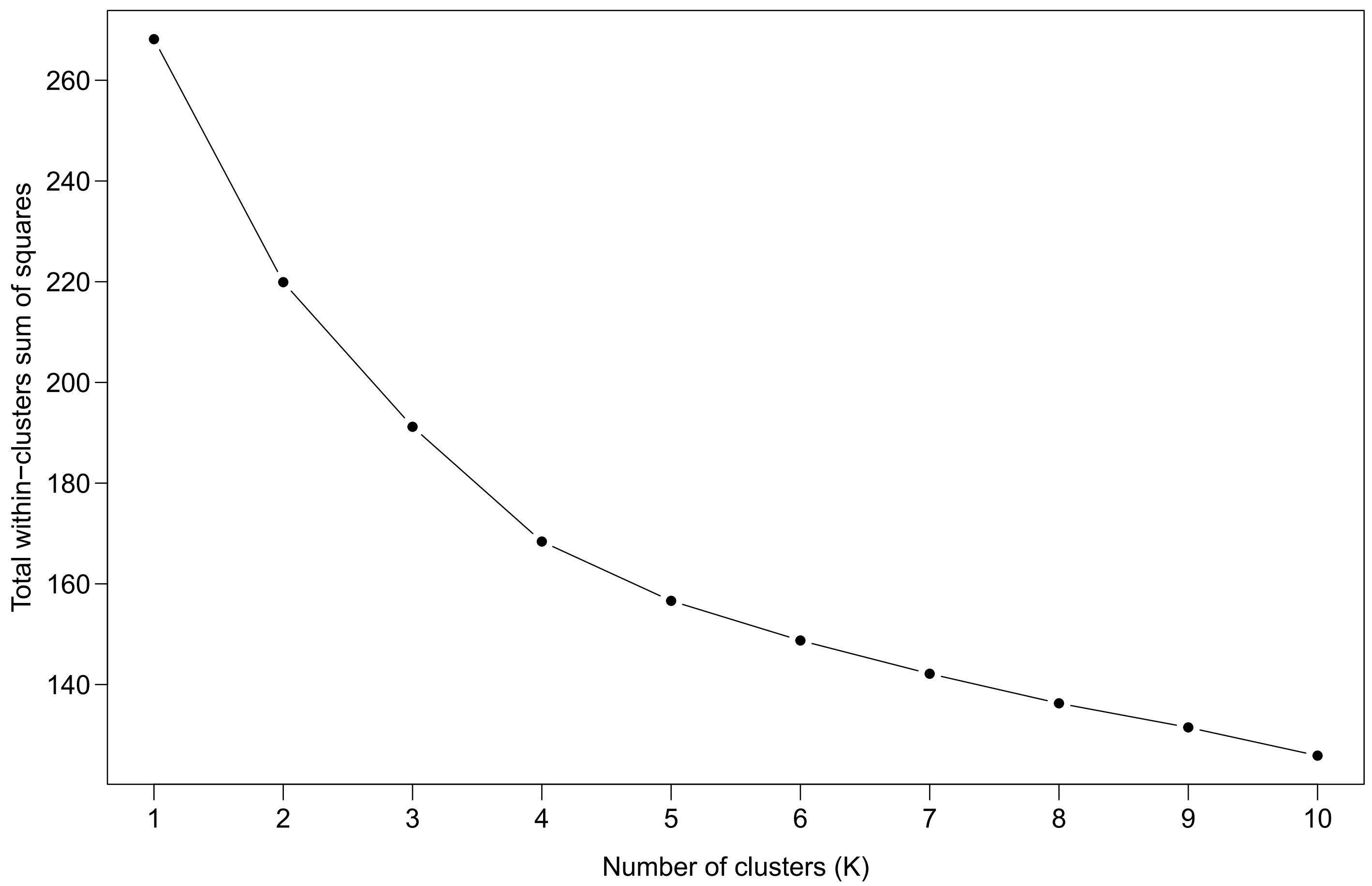


**Fig. S1. The optimal number of clusters (K) using the elbow method.** The point where the rate slows for the total within-clusters sum of squares is the optimal number of clusters derived from the hierarchical cluster analysis.

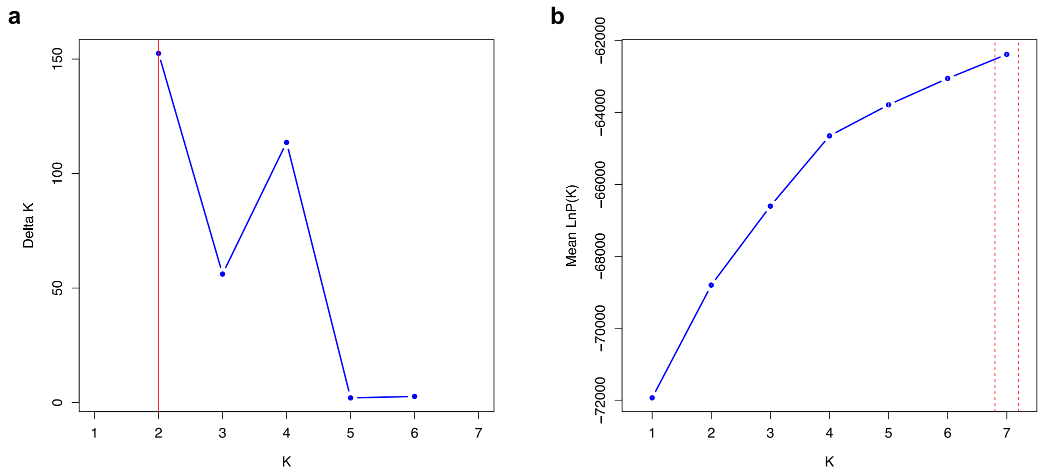


**Fig. S2. STRUCTURE best K values.** STRUCTURE best K values for K=1-7 based on **a)** ΔK (the second order rate of change in the log probability of data between successive K values), and **b)** mean LnP(K) (the mean log probability of the data). Red lines indicate the optimal K values.

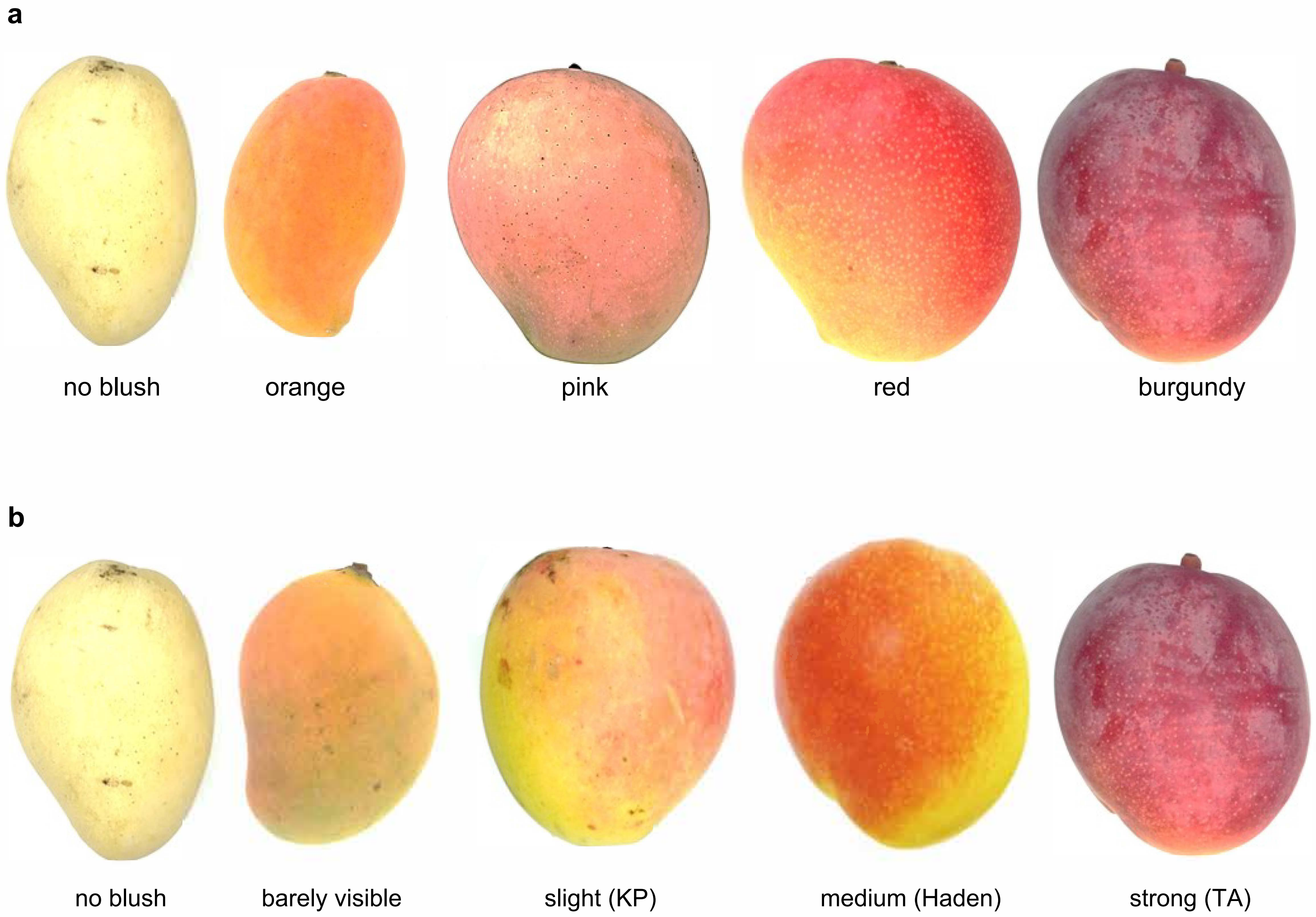


**Fig. S3. Fruit trait categories. a)** Fruit blush colour is split into five categories. **b)** Fruit blush intensity increases from no blush to strong blush on an ordinal scale. The cultivars in brackets best reflect the colour intensity, where KP is Kensington Pride and TA is Tommy Atkins.

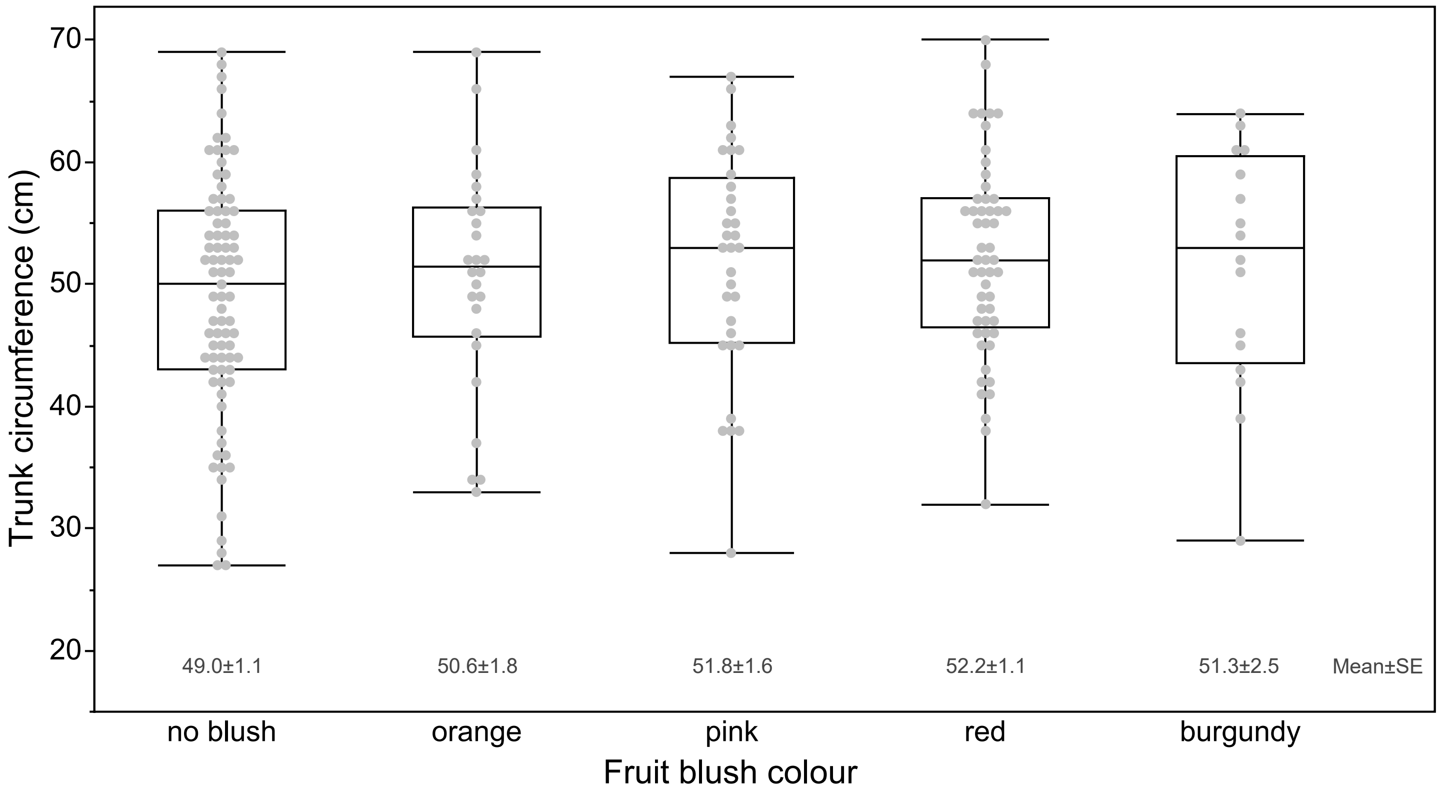


**Fig. S4. The mean and standard error (SE) of trunk circumference across fruit blush colour categories.**

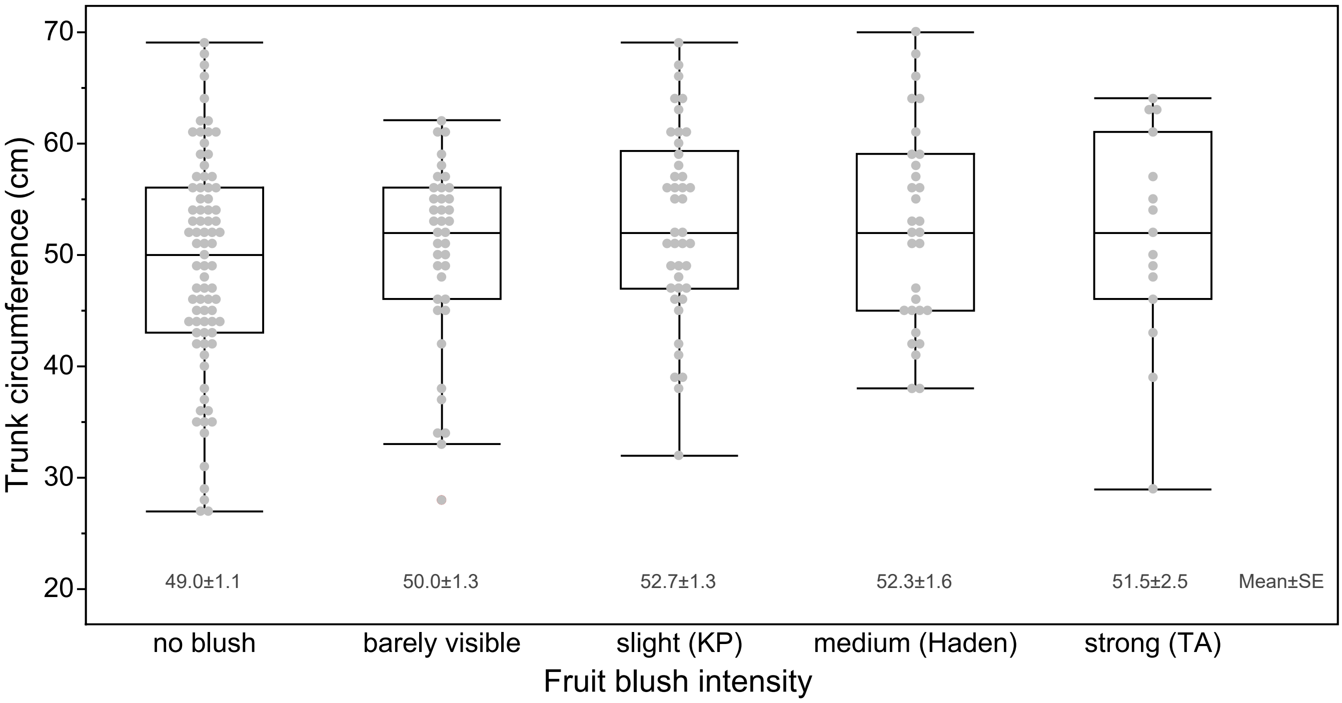


**Fig. S5. The mean and standard error (SE) of trunk circumference across fruit blush intensities.** The cultivars in brackets best reflect the colour intensity, where KP is Kensington Pride and TA is Tommy Atkins.
